# Let’s talk about sex: Male and female mice show similar fear memory retention despite hippocampal activity differences during encoding and consolidation

**DOI:** 10.64898/2026.01.28.702335

**Authors:** Katherine O. McDonald, Temmie Yu, Aditi Prabhu, Sara J Aton

## Abstract

Accurate and efficient memory processing is essential for survival. A body of ongoing work in both human subjects and animal models suggests that memory processing may differ substantially between males and females. In mice, contextual fear memory (CFM) encoding, consolidation, and recall have been well studied, and the mouse hippocampus and amygdala have been implicated in these processes. The present study addresses whether the activation of these brain regions differs between male and female mice at each stage of CFM processing. We find that male and female C57BL6 mice show no differences in sleep behavior following single-trial contextual fear conditioning (CFC), which is essential for CFM consolidation. We also find no significant differences in CFM recall behavior between male and female mice. However, females - but not males - show significantly increased expression of cFos in dorsal hippocampal CA1 and CA2 neurons during CFM encoding. On the other hand, males – but not females - show increased cFos expression among dentate gyrus (DG) granule cells, and decreased cFos expression within dorsal CA1 and CA3 during CFM consolidation. Males also show significant decreases in dorsal CA1 cFos expression during memory recall. These findings highlight the idea that the neurobiological underpinnings of memory processing may differ between males and females, even when recall performance is identical.

**Highlights:** Historically, research on the neurobiological basis of memory processing has been carried out mainly in male subjects. Thus, our understanding of these mechanisms is biased towards male brain neurophysiology. This is problematic, as numerous studies have reported sex-based performance differences for episodic memory tasks, in which male subjects have variously performed better, worse, or the same as females. Here, we find that male and female mice perform similarly on a well-studied experimental memory task but nonetheless have differences in the relative activity of different brain structures during sequential stages of memory processing. This emphasizes the importance of including both males and females in memory studies, due to potential sex differences in the neurobiological substrates of memory.

## Introduction

Over the past 50 years, our understanding of biological processes underlying memory processing - an understanding essential for preserving healthy cognition - has advanced substantially. However, many of the cellular, molecular, and systems-level mechanisms of memory processing have been described in studies using only male animals ^1^. This is surprising, as many studies have found substantial differences in memory processing mechanisms between males and females ^2^. Closing this knowledge gap has important implications. For example, men and women differ in their rate of Alzheimer’s disease diagnosis across all age groups^3,4^, and the genetics associated with cognitive resilience and susceptibility to dementia can also differ between males and females ^5,6^. Understanding sex differences in the memory processing mechanisms may allow more tailored interventions for Alzheimer’s disease and other disorders characterized by disrupted cognition. The exclusion of female animals from studies addressing memory mechanisms severely limits our ability to extrapolate experimental findings to the human population.

Episodic memories - memories of events, associations, and contexts - are disrupted in many neurological disorders and rely heavily on hippocampal and neocortical regions for proper encoding, consolidation, and recall ^7–10^. Contextual fear memory (CFM) is among the best-studied episodic-like memory used in laboratory mice, where it can be encoded in a single learning trial (referred to as contextual fear conditioning [CFC]). CFC itself activates the amygdala and hippocampus ^11,12^, and CFM is consolidated in the hours following CFC via a mechanism that relies on dorsal hippocampal neural activity ^13^, gene transcription ^14,15^, and protein translation ^16–18^. While the process of CFM consolidation is not fully understood, post-CFC sleep appears to play an important role via its effects on hippocampal and neocortical activity patterns ^19–22^ and its activation of critical intracellular pathways ^15,16,23–26^. More recent work has shown that CFM recall evokes reactivation of the same neurons activated during CFC - including populations of so-called “engram” neurons in the hippocampus and amygdala ^15,27^.

Historically, most research on CFM has been carried out using only male animals, excluding females with the aim of reducing potential variability in behavior or physiology across the estrus cycle ^1,28^. A few recent studies have begun to test sex as a biological variable affecting CFM processing. Some of these comparative studies have shown that male rodents show either greater contextual freezing responses, or more specific CFM responses than females (i.e., with less context generalization) during CFM recall ^29–31^. However, others identified either no behavioral differences between males and females (and even similar behavioral variability between males and cycling females ^2832^), or greater freezing responses in females ^29^. The mechanistic underpinnings of these putative sex-based CFM differences are also unclear. For example, CFM consolidation is dependent on post-CFC sleep ^15,19,23,25,33^, and differences in sleep architecture between male and female rodents have been reported ^34–36^. It is unknown whether these sleep architecture differences could translate to differences in CFM consolidation between males and females. Intriguingly, even in studies where CFM recall appears equal between males and females, the cellular pathways required for memory processing may vary with sex ^37–39^. In the context of these varying findings, it remains unclear whether the activity of circuits engaged by CFM processing also differs between males and females ^2,40^. Our present study investigates potential sex differences in activity of hippocampal and amygdalar structures during CFM encoding, consolidation, and recall, in the absence of a behavioral effect. Our data highlight the need for further investigation of different brain regions’ contribution to memory processing in males vs. females.

## Methods

### Mouse husbandry and housing

Prior to each experiment, C57BL6/J male (*n* = 17) and female (*n* = 18) mice (Jackson; 3-6 months of age) were maintained on a 12 h light/12 h dark cycle (lights on at 9 am) under controlled temperature and relative humidity conditions (22 ± 2°C; 60-75%). Mice were group-housed in standard filter-top caging with age-matched littermates (no more than 5 to a cage), provided beneficial enrichment (Enviro-Dry nesting packets), and allowed *ad lib* access to food and water. Beginning on the first day of habituation to experimental handling, each mouse was single-housed with additional enrichment (cotton nestlets) to decrease effects of cage mate interactions on post-learning sleep, and remained single-housed for the duration of behavioral experiments (4-5 days total). All animal husbandry and experimental procedures were approved by the University of Michigan Institutional Animal Care and Use Committee (IACUC). Because behavioral outcomes in mice can vary with sex of the experimenter ^41^, all handling procedures were carried out by a female experimenter. All mice were monitored daily to ensure their health during experiments.

### Contextual fear conditioning (CFC) and contextual fear memory (CFM) testing

Prior to each experiment, all mice were handled daily for 3 days (4 min/day) for habituation to experimenter handling procedures. Mice were assigned to experimental timepoint groups for measuring activity-driven protein expression during CFM encoding (*n* = 4 male, 4 female), consolidation (*n* = 4 male, 4 female), or recall (*n* = 4 male, 4 female) (**Fig. 1A**). A group of age-matched control mice (*n* = 5 male, 6 female) underwent the same housing and handling procedures but did not undergo CFC. The number of mice per group was estimated to be sufficient for detecting sex differences in our primary outcome measure (immunohistochemical measures), based on prior studies^29^. To minimize batch effects, control and experimental mice of both sexes underwent behavioral procedures simultaneously. Mice were assigned to groups quasi-randomly, distributing littermates between experimental groups so that: 1) no two mice in a single group came from the same litter and 2) there was an even age distribution across groups. On the fourth day, experimental mice underwent single-trial CFC as previously described ^15^. At lights on (Zeitgeber Time [ZT] 0), each mouse was placed in a novel cylindrical plexiglass conditioning chamber (22 cm diameter × 25 cm high) with a black and white checkerboard pattern on the interior walls, a metal grid floor for foot shock delivery (Med Associates), and an associated scent cue (lemon Lysol). Mice were allowed to explore this environmental context for 2 min 28 s before receiving a 2-s, 0.75 mA foot shock, followed by an additional 30 s in the chamber, and were then returned to their home cage.

**Figure 1:**
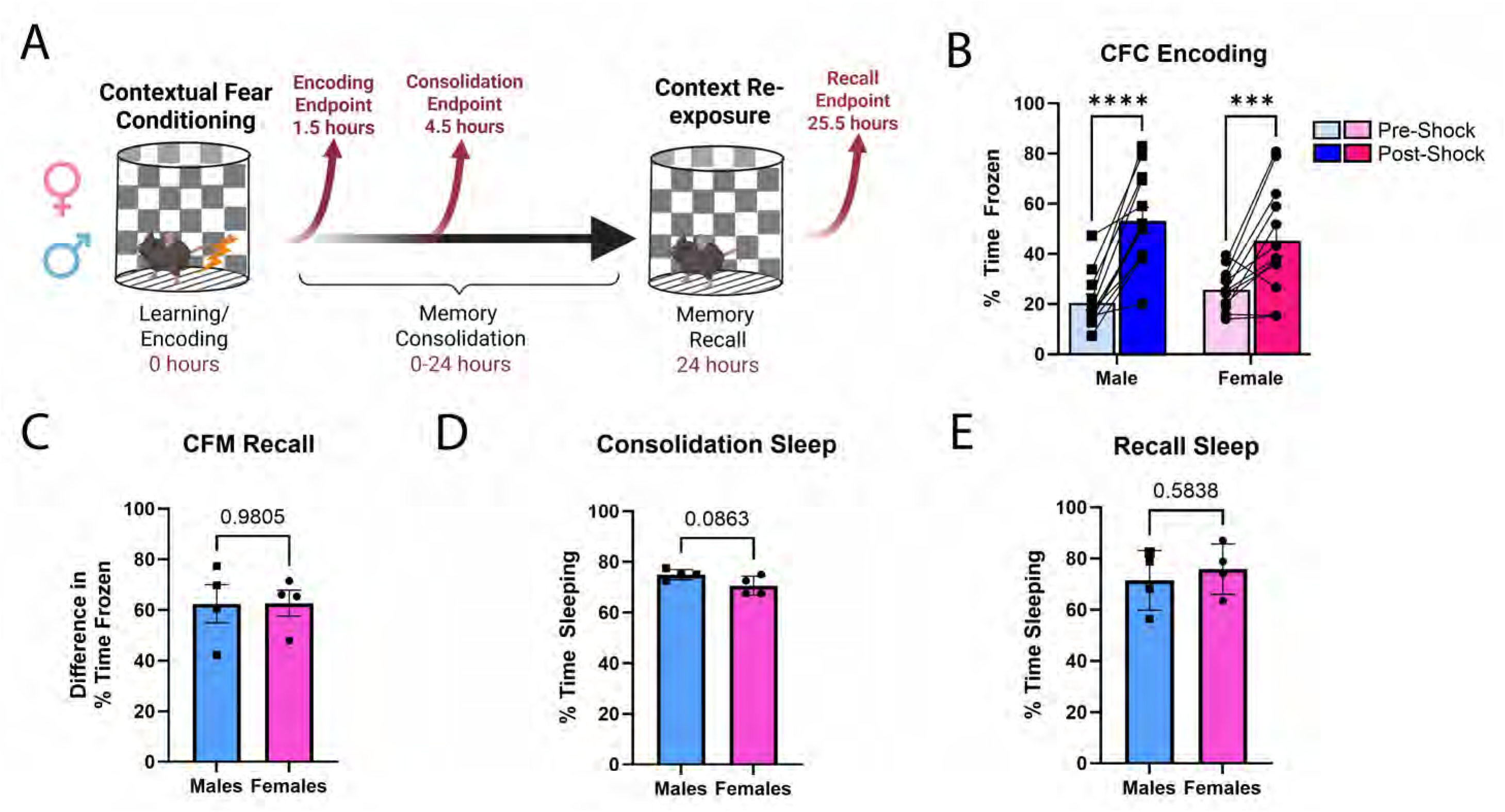
Male and female mice show similar behavioral responses to CFC, post-CFC sleep, and CFM recall. **A)** Male and female experimental mice underwent single-trial CFC, then were euthanized for immunohistochemical assessment of neural activity patterns during CFM encoding, consolidation, or recall. **B)** During single-trial CFC, both male and female mice showed significantly greater freezing post-shock compared to pre-shock (*n* = 12 male, 12 female, two-way ANOVA with uncorrected Fisher’s LSD males post-shock vs. pre-shock *p* < 0.0001, females post to pre-shock *p* = 0.0008, other comparisons *p >* 0.05, main effect of sex F[1, 22] = 0.082, *p* = 0.78, η^2^ = 0.0014; main effect of memory stage F[1,22] = 55.69, *p* < 0.0001, η^2^ = 0.42; sex × memory stage interaction effect F[1, 22] = 3.9, *p* = 0.061, η^2^ = 0.029). **C)** Male and female mice showed similar CFM recall – measured as % change in context-specific freezing - during testing 24 h following CFC (*n* = 4 males, 4 females, male vs. female *p* = 0.98, unpaired Student’s t test, t = 0.026, df = 6, R^2^ = 0.00011). **D)** Male and female animals from the CFM consolidation group spent similar % time asleep in the 4.5 hours following CFC (males mean ± SEM = 75% ± 1.021%, female average mean ± SEM = 70.63% ± 1.88%, *n* = 4 males 4 females, male vs female *p* = 0.086, unpaired Student’s t test t = 2.05, df = 6, R^2^ = 0.42). **E)** Male and female animals from the CFM recall group also spent similar % time sleeping over the first 6 h following CFC (males mean ± SEM = 71.47% ± 5.83%, female average mean ± SEM = 75.88% ± 4.91%, *n* = 4 males 4 females, male vs female *p* = 0.58, unpaired Student’s t test t = 0.58, df = 6, R^2^ = 0.053). All values indicate mean ± SEM.

Timepoints for euthanasia and brain tissue harvest were selected based on prior studies indicating that cFos protein expression peaks roughly 1.5 h following experimentally-induced neuronal activity ^42–45^. 1.5 h after CFC (a timepoint at which cFos protein expression would be expected to reflect neuronal activity associated with CFM encoding) mice in the encoding experimental group (*n* = 4 male, 4 female) and 3 time-matched, no-CFC control mice (*n* = 1 male, 2 female) were euthanized with an i.p injection of pentobarbital (Euthasol; Butler Schein) prior to transcardial perfusion with 15 ml of ice cold phosphate buffered saline (PBS) followed by 15 ml of ice cold 4% paraformaldehyde. Mice in the consolidation experimental group (*n* = 4 male, 4 female) were euthanized and perfused 4.5 h after CFC (to quantify neural cFos expression associated with active CFM consolidation)^15,25^ along with 4 time-matched control mice (*n* = 2 male, 2 female). The 4.5-h post-CFC timepoint corresponded to cFos protein expression driven by brain activity after 3 h of post-learning sleep, a time period during which cellular pathways associated with memory consolidation mechanisms are upregulated in dorsal hippocampus ^25,26,46^. The next day at lights on, mice in the recall experimental group (*n* = 4 male, 4 female) were re-exposed to the CFC context for 5 min for assessment of CFM recall and were then returned to their home cage. These mice and 4 no-CFC control animals (*n* = 2 male, 2 female) were euthanized and perfused 1.5 h after CFM recall to capture neuronal activity associated with recall. Behavior during CFC and CFM recall was video monitored. Freezing behavior during both recordings was quantified using an automated analysis algorithm in EthoVision software by a blinded researcher. For each mouse, CFM-associated freezing behavior was expressed as (% of time spent freezing during the 5-min recall session) - (% of time freezing in the 1 min 28 s period prior to CFC foot shock) to account for individual differences in baseline freezing as done previously ^19–21^. All fear conditioned mice included in the analysis displayed a characteristic behavioral response to foot shock, including post-shock freezing. Experimental data for a single male assigned to the consolidation-timepoint control group was removed from subsequent analysis due to being incorrectly sexed at the time of weaning.

### Sleep monitoring

All mice were allowed *ad lib* sleep in their home cages, with visual sleep monitoring starting either immediately after CFC (for experimental animals) or at lights on (for control animals). Sleep behavior was identified visually based upon mice becoming quiescent, nesting, and assuming typical sleep postures (crouched within the nest with head down, motionless outside of regular respiratory movements). This non-invasive method for identifying sleep behavior is comparable to polysomnographic (EEG [electroencephalogram] and EMG [electromyography]) recording ^47^, and can be carried out without prior surgical procedures and minimal prior single housing. Each mouse in the consolidation and recall groups was observed continuously over the first 4.5-6 h following CFC (4.5 h between CFC and sacrifice for mice in the consolidation group, and the first 6 h post-cfc for the recall group). The first few hours following CFC is a window of time during which uninterrupted sleep is critical for successful CFM consolidation ^15,19,23,25,33^. Mice were determined to be either asleep or awake once every 5 min, in real time, by a single researcher blinded to experimental condition, across this 4.5-6-h time window. For each mouse, sleep behavior was quantified as a % of the total observation period.

### Estrus cycle monitoring

Because vaginal smear collection for cytological assessment can cause distress and disrupt cognition in female mice ^48^, assessment of estrus cycle stage in females was carried out during tissue harvest by imaging the anogenital area for visual assessment of the exterior vaginal canal, using established methods ^49^. Distributions of estrus cycle phases for females in each experimental group is shown in **Supplementary Table 1**. The estrus cycle could not be confidently determined in two female animals, and thus their cycle stage is not reported.

### Immunohistochemistry and image analysis

Immediately following perfusion, brains were extracted, incubated in 4% paraformaldehyde at 4°C for 24 h, then transferred to ice cold phosphate-buffered saline (PBS) for coronal sectioning (80 µm thickness) using a vibratome (Leica VT1200s). Coronal sections approximately -1.58 to -2.18 mm from bregma were selected for immunostaining. Sections were blocked in PBS with 1% Triton X-100 and 5% normal donkey serum for 24 hours at 4°C, incubated for 4 days at 4°C in primary antibody (rabbit anti cFos 1:750 [Abcam; ab190-289]), washed in 0.2% Triton X-100 in PBS, and incubated for 2 days at 4°C with secondary antibody (DyLight 633 donkey anti rabbit igG 1:800 [Novus Biologicals; NBP1-75638]). Antibody dilutions and incubation periods were based on prior studies of neuronal activation in mouse hippocampus following CFC ^15,50^. After immunostaining, brain sections were washed in PBS, incubated at room temperature in DAPI (1:500) for 5 min to stain cell nuclei, re-washed, and mounted and coverslipped with ProLong Gold antifade reagent (ThermoFisher; P36930). All tissue samples were prepared simultaneously. Immunolabeled brain sections were imaged on a Leica SP8 laser scanning confocal microscope at 40× magnification. For each brain region quantified, 4-5 serial sections were imaged for each animal. Quantifications for each of these sections were averaged, to obtain a single numeric value for each region.

Brain structures within acquired images were mapped in reference to the Allen Brain Reference Atlas, using QuPath image analysis software. For all hippocampal subregions (CA1-CA3 and dentate gyrus [DG]), analysis of cFos expression was constrained to the principal cell body layers. To ensure that CA1-CA3 measurements were fully contained within the appropriate subregions (i.e., where anatomical borders between these regions are not clearly defined), analysis was constrained to the center of each region (approximately 100 µm from borders of neighboring regions of interest), avoiding areas close to the border with other CA structures. Individual cells were detected within each brain region using DAPI fluorescence. For basolateral amygdala (BLA), lateral amygdala (LA), and dorsal CA1-3 regions, QuPath software was used for intensity threshold-based detection of cFos+ cells; automated detection of cFos+ cells was curated by an experimenter blinded to the sex and experimental conditions of each mouse. A DAPI+ identified nucleus was considered cFos+ if the average pixel intensity within its boundaries was > 175 (on a scale from 0 to 255). cFos+ ratios were calculated as the number of cFos+ cells over the total number of DAPI+ cells per region. High nuclear density within the DG granule cell body layer precluded accurate DAPI+ cell detection. To accommodate this technical issue, cFos+ cells were manually counted across the granule cell body layer by an investigator blinded to experimental conditions and reported as cFos+ cells/mm^2^ for the DG only. Scoring reliability was verified by a second blinded investigator in a subset of samples (linear regression analysis of inter-rater scores for animal samples R^2^ = 0.60 *p* =0.0091, data not shown).

For all image analyses, values for experimental mice (male or female encoding, consolidation, or recall) were compared to a group of control mice (*n* = 5 male, 6 female) that did not undergo CFC and were pooled across time points. Sample collection time point did not affect cFos expression levels among control mice.

### Statistical Analysis

For all analyses comparing two groups (**Fig. 1C-E**), simple unpaired t tests were used for statistical analyses. For all comparisons of more than two groups (**Fig. 1B** and **Figs. 2**–**6**), two-way ANOVA were performed. All two-way ANOVA analyses compared experimental group means (encoding, consolidation, and recall) of both sexes (male and female) to male and female control group means. Normality of residuals for all analyses were assessed with the Shapiro-Wilk test and passed normality assessment (alpha = 0.05). Datasets that were not normally distributed (**Fig. 3B** and **Fig. 4B**) were log-transformed for analysis with two-way ANOVA. All transformed datasets passed the Shapiro-Wilk normality test. Heteroscedasticity was assessed using the Spearman’s rank correlation test for two-way ANOVAs or using the F test for unpaired t tests. Datasets that did not pass the heteroscedasticity test were log transformed, and reported statistics run on transformed data (**Fig. 2B** and **2D**, **Fig. 4B**). All t tests passed the F test to compare variances. All transformed datasets are displayed in their raw form, with appropriate statistical markers for clarity. For analyses that compared the same animal over multiple time points (**Fig. 1B**), a two-way ANOVA with uncorrected Fisher’s LSD was performed. For all other two-way ANOVAs, Dunnett’s *post hoc* test was performed to correct for multiple comparisons. Effect sizes are reported as R^2^ for all unpaired t tests, and η2 for all two-way ANOVAs. All data analyzed using GraphPad Prism 11.0.0.

**Figure 2:**
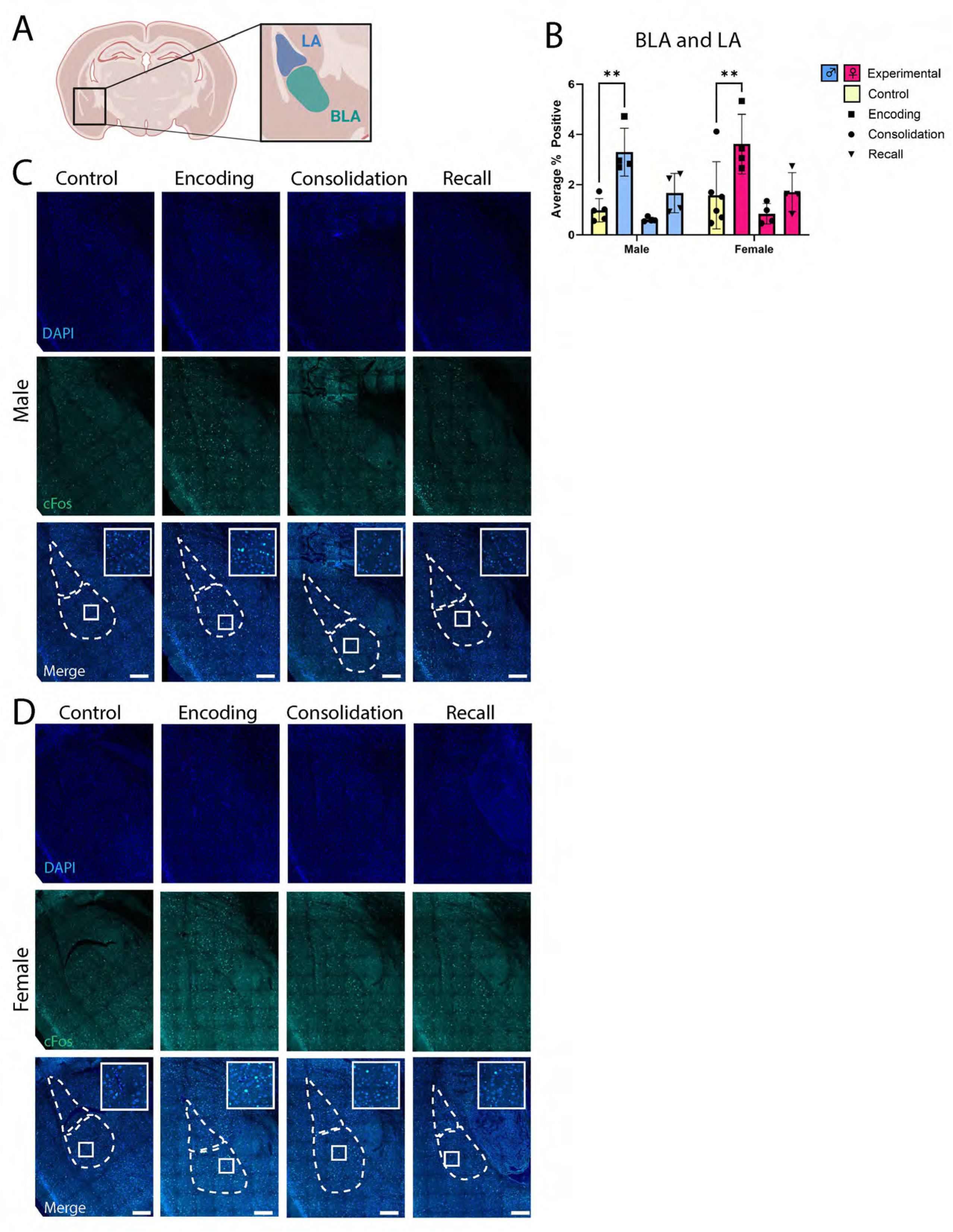
Amygdalar neuron activation during CFM encoding is similar between male and female mice. **A)** The basolateral and lateral amygdala (BLA and LA) were immunolabeled to compare the density of cFos+ neurons across each stage of CFM processing (encoding, consolidation, and recall) in male vs. female mice. **B)** In both males and females, the proportion of cFos+ BLA and LA neurons was significantly greater during CFM encoding compared to control mice. There were no significant differences in consolidation and recall groups compared to controls for either sex (two-way ANOVA with Dunnett’s *post hoc* test, male encoding vs. control *p* = 0.0018, female *p* = 0.0074 all other comparisons *p* > 0.05. *n* = 5 male, 6 female control; 4 male, 4 female encoding; 4 male, 4 female consolidation; 4 male, 4 female recall, main effect of sex F [1,27] = 0.55 *p* = 0.47, η^2^ = 0.00043; main effect of memory stage F [3, 27] = 25.46, *p* < 0.0001, η^2^ = 0.25; sex × memory stage interaction effect F [3, 27] = 0.032, *p* = 0.99, η^2^ = 0.00044). **D)** Representative images of male and **E)** female amygdalar structures with DAPI+ and cFos+ immunolabeling. All values indicate mean ± SEM. Scale bars = 250 μm.

**Figure 3:**
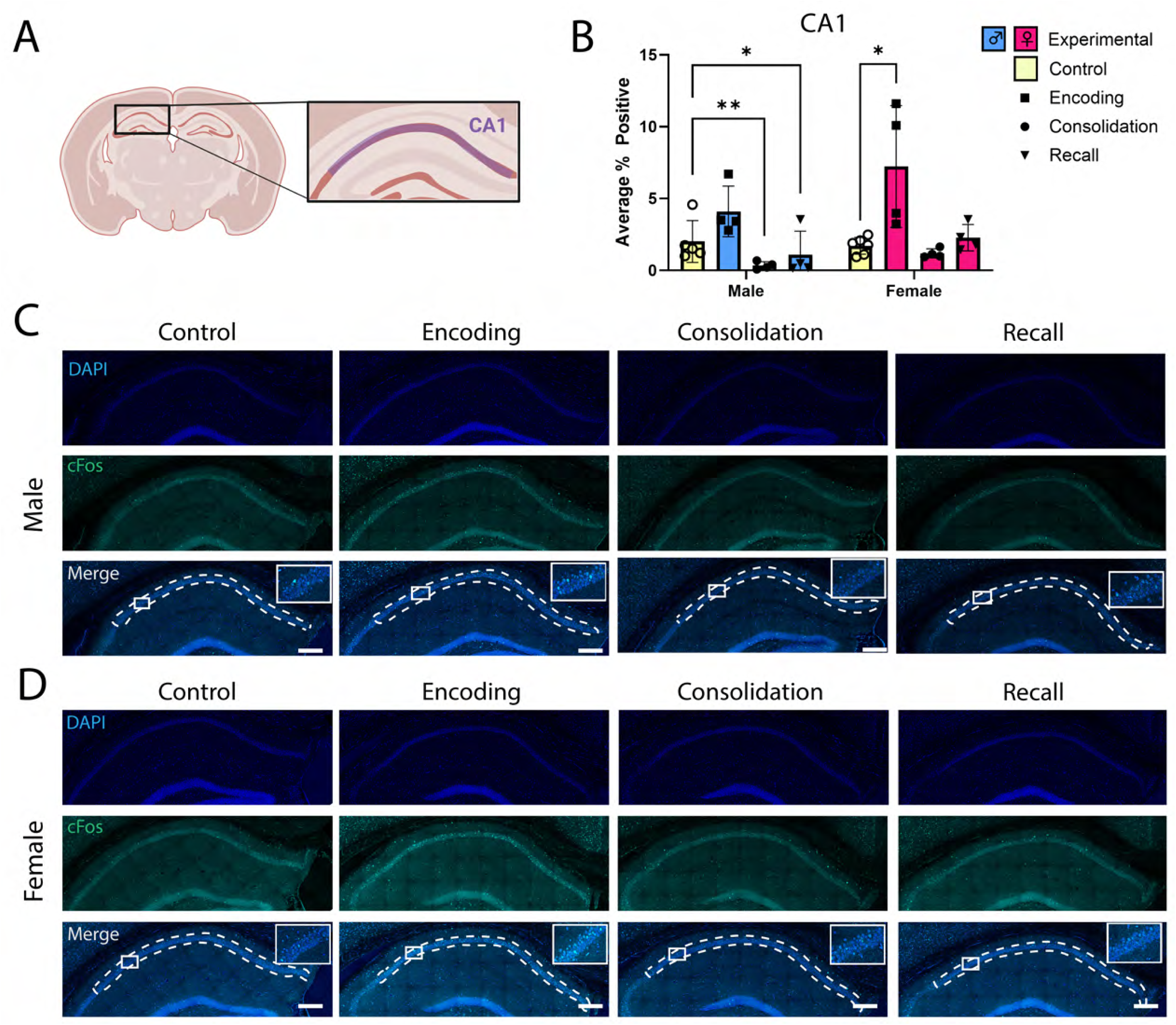
CA1 pyramidal neuron activity differs between male and female mice across CFM processing stages. **A)** The proportion of cFos+ neurons in the dorsal CA1 pyramidal cell body layer was quantified for male and female mice in each group. **B)** Female mice - but not male mice - had a significantly greater proportion of cFos+ CA1 neurons after CFM encoding compared to controls (two- way ANOVA with Dunnett’s *post hoc* test female encoding vs. control *p* = 0.016). Additionally, male mice - but not female mice - showed a reduction in cFos+ CA1 neurons during CFM consolidation (two- way ANOVA with Dunnett’s *post hoc* test male consolidation vs. control *p* = 0.0014) and recall compared to control (male recall vs control *p* = 0.027, all other comparisons *p* > 0.05), with significant main effects of both sex and memory stage (*n* = 5 male, 6 female control; 4 male, 4 female encoding; 4 male, 4 female consolidation; 4 male, 4 female recall. Main effect of sex F [1, 27] = 12.75, *p* = 0.0014, η^2^ = 0.14; main effect of memory stage F [3, 27] = 14.16, *p* < 0.0001, η^2^ = 0.48; sex × memory stage interaction effect F [3, 27] = 2.83, *p* = 0.057, η^2^ = 0.096). **C-D)** Representative CA1 images with DAPI and cFos immunolabeling, for **C)** male and **D)** female mice in each group. Values indicate mean ± SEM. Scale bars = 250 μm.

**Figure 4:**
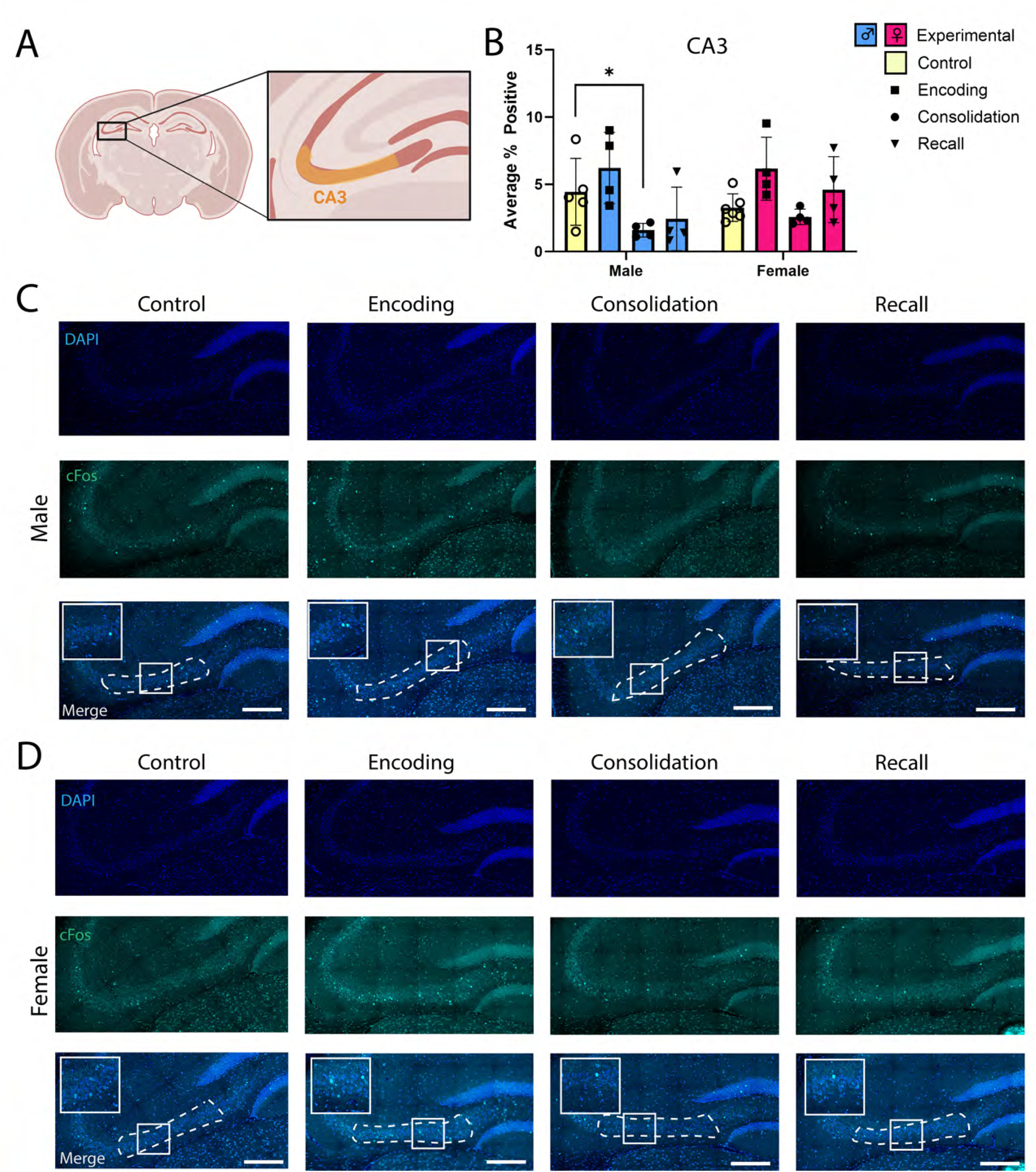
CA3 pyramidal neuron activity is suppressed during CFM consolidation in male mice. **A)** The proportion of cFos+ dorsal CA3 pyramidal neurons was compared male and female mice at each CFM processing stage. **B)** Male mice had significantly lower cFos+ neuron density during consolidation compared to controls (two-way ANOVA with Dunnett’s *post hoc* test male consolidation vs. control, *p* = 0.024), while other groups were not significantly different from controls (all other comparisons, *p* > 0.05), with a significant overall main effect of memory stage *n* = 5 male, 6 female control; 4 male, 4 female encoding; 4 male, 4 female consolidation; 4 male, 4 female recall. Main effect of sex: F [1, 27] = 2.9, *p* = 0.099, η^2^ = 0.052; main effect of memory stage: F [3, 27] = 7.05, *p* = 0.0012, η^2^ = 0.38; sex × memory stage interaction: F [3, 27] = 2.031, *p* = 0.13; η^2^ = 0.11). **C-D)** Representative CA3 images from **C)** male and **D)** female mice in each group. Values indicate mean ± SEM. Scale bars = 250 μm.

## Results

### Males and females show comparable CFM recall and post-CFC sleep behavior

We first compared learning-related behavior during single-trial CFC, as well as during subsequent CFM recall between male and female mice (**Fig. 1A**). Males and females did not differ in the amount of time spent in freezing behavior during CFC itself (i.e., during CFM encoding) (**Fig 1B**; males pre shock mean ± SEM = 20.52 ± 3.17%, males post shock mean ± SEM = 54.21 ± 5.43%, females pre shock mean ± SEM = 26.01 ± 2.3%, females post shock mean ± SEM = 45.59 ± 6.53%, main effect of sex: F[1, 22] = 0.082, *p* = 0.78, main effect of memory stage: F[1,22] = 55.69, *p* < 0.0001, sex × memory stage interaction: F[1, 22] = 3.9, p=0.061). Both male and female mice showed increased freezing behavior upon return to the CFM context (**Fig 1B-C**; 62.5 ± 7.6% and 62.7 ± 5.1% [mean ± SEM] increases in freezing for males and females, respectively), and CFM recall did not differ between males and females (*p* = 0.98, unpaired t test;) (**Fig. 1B**). Because episodic memory consolidation is highly dependent on sleep ^15,19,23–25,33,51^, mice from consolidation and recall groups underwent visual scoring of sleep behavior over the first 6 h following CFC. There were no observed significant differences between male and female mice with respect to sleep behavior during this time window (consolidation group males and females: 75.0 ± 1.02% and 70.63 ± 1.88% [mean ± SEM] of 6 h post-CFC spent asleep, respectively, *p* = 0.086 unpaired t test; recall group males and females 71.47 ± 5.83% and 75.88 ± 4.91% [mean ± SEM] time spent asleep, respectively, *p* = 0.58 unpaired t test) (**Fig. 1C-D**).

### Amygdalar activity increases in response to fear learning in both males and females

Fear memory processing has been shown to activate neurons in amygdalar structures, including the lateral (LA) and basolateral amygdala (BLA) ^27,52^. To test whether sex differences in amygdalar activation were present in the absence of a behavioral difference in CFM recall, we quantified the proportion of amygdalar neurons that were cFos+ during CFM encoding, consolidation, and recall stages. The two amygdalar structures (BLA and LA) were quantified together (**Fig. 2A**), although separate quantification of the two regions is shown in **Supplementary Fig. 2**. We compared the proportion of amygdalar cells expressing cFos protein in males vs. females during each stage of CFM processing, identifying changes in mice undergoing CFC vs. home cage control mice. Male and female mice showed a similar degree of amygdalar activation immediately following CFC, with a significant increase in cFos+ cell proportion during encoding compared to controls, which returned to control levels during CFM consolidation (**Fig. 2B-C**).

### Hippocampal CA1-CA3 activity patterns differ between males and females across CFM encoding, consolidation, and recall

Neuronal activity in dorsal hippocampal regions CA1 and CA3 is necessary for successful CFM processing ^13^. To test for potential sex differences in dorsal hippocampus during CFM processing, we next compared the proportion of CA1 principal cells that were cFos+ in males and females across encoding, consolidation, and recall (**Fig. 3A-C**). Our data indicate that in the CA1 of females - but not males - activity during encoding is significantly elevated compared to control animals (**Fig. 3B**). Interestingly, the proportion of cFos+ CA1 neurons was significantly *decreased* during both CFM consolidation and recall in male mice - but not female mice - compared to controls.

In contrast to CA1, CA3 (**Fig. 4A-C**) did not show a significant increase in cFos+ neuron density during CFM encoding in either male or female mice. Female mice - but not male mice - showed a trend for increased proportion of cFos+ CA3 neurons during encoding (two-way ANOVA with Dunnett’s *post hoc* test, male encoding vs. control *p* = 0.49, female encoding vs. control *p* = 0.15). In contrast, male mice - but not female mice - showed a significant *reduction* in cFos+ neuron density in CA3 during memory consolidation (Dunnett’s *post hoc* test, male consolidation vs. control *p* = 0.023; **Fig. 4D**).

While the roles of CA1 and CA3 in CFM processing have been well studied, contributions of adjacent CA2 are less clear ^53^. Similarly to our findings in CA1, CA2 (**Fig. 5A-C**) had significantly more cFos+ cells during CFM encoding compared to controls in female - but not male - mice (two-way ANOVA with Dunnett’s *post hoc* test, *p* = 0.015). Unlike the CA1, no significant changes in CA2 cFos expression were observed in either sex during the consolidation or recall of CFM (**Fig. 5D**).

**Figure 5:**
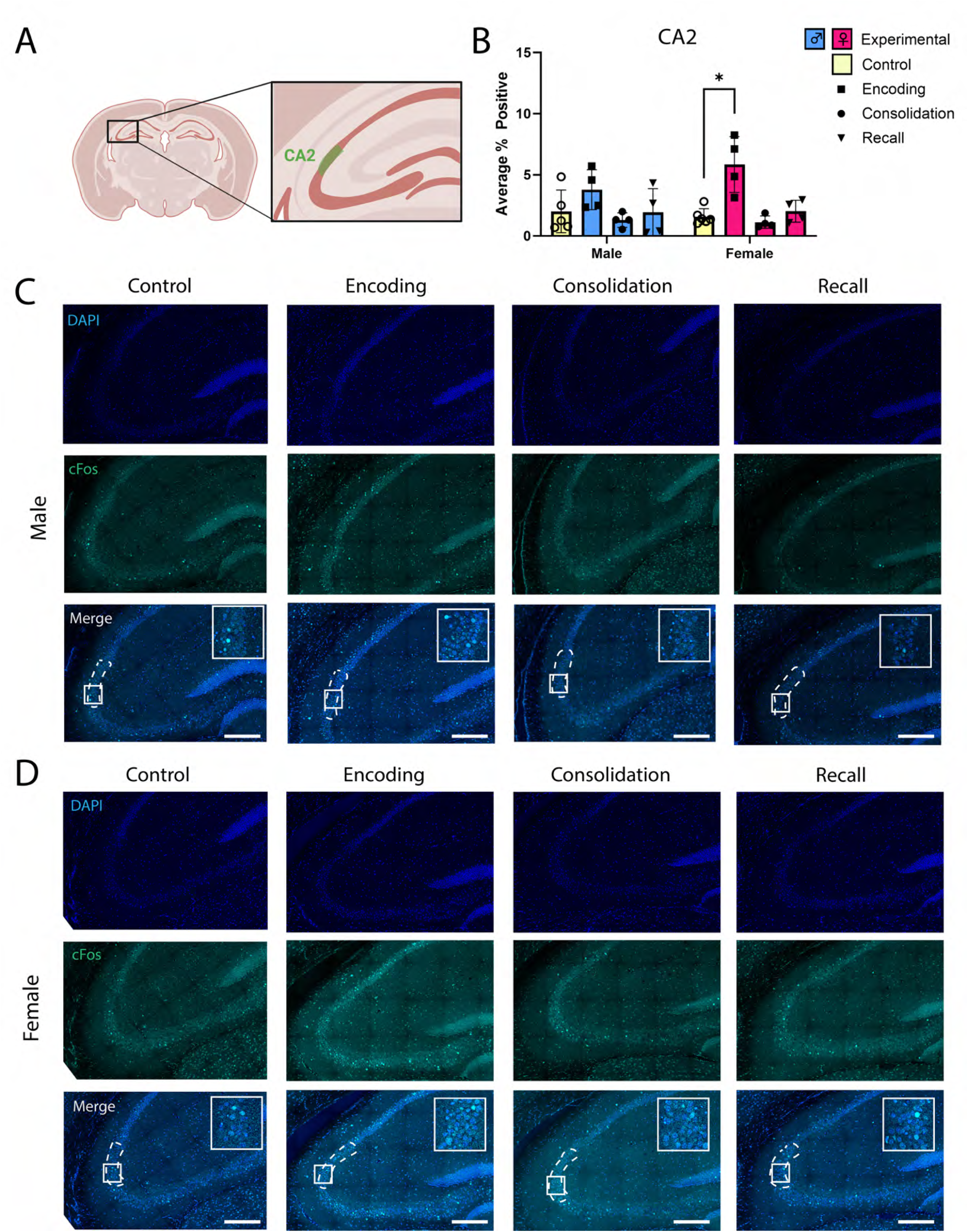
CA2 pyramidal neuron activity is significantly increased during CFM encoding in female mice only. **A)** cFos+ neuron density was quantified in the pyramidal cell body layer of dorsal CA2. **B)** Female mice had significantly greater CA2 cFos+ neuron density during encoding compared to controls (two-way ANOVA with Dunnett’s *post hoc* test female encoding vs. controls *p* = 0.0153), while the same was not true for the male cohort (all other comparisons *p* > 0.05), with a significant overall main effect of memory stage (*n* = 5 male, 6 female control; 4 male, 4 female encoding; 4 male, 4 female consolidation; 4 male, 4 female recall. Main effect of sex F [1, 27] = 0.9250, *p* = 0.6847, η^2^ = 0.019; main effect of memory stage F [3, 27] = 6.754, *p* = 0.0015, η^2^ = 0.41; sex × memory stage interaction effect F [3, 27] = 0.50, *p* = 0.69; η^2^ = 0.030. **C-D)** Representative CA2 images from **C)** male and **D)** female mice in each group. Values indicate mean ± SEM. Scale bars = 250 μm.

### Dentate gyrus (DG) activity is increased during active CFM consolidation in male mice only

The hippocampal DG is another structure engaged by CFC encoding and recall, which more recently has been implicated in CFM consolidation ^15,46,50^. However, aside from a recent report of greater DG activity during CFC in males vs. females ^29^, it is largely unknown whether sex differences in CFM processing-associated activity exist within this region. While in hippocampal CA1-CA3, cFos expression was highest during CFC and no increase was observed during CFM consolidation (**Figs. 3**–**5**), no increase in DG cFos expression was observed during CFC in either males or females (**Fig. 6A-C**). In contrast, during CFM consolidation, the proportion of cFos+ DG neurons increased in male mice (**Fig 6D**; two-way ANOVA with Dunnett’s *post-hoc* test, *p* = 0.0011) with a significant sex × memory stage interaction effect (F [3, 27] = 3.39, *p* = 0.03). Surprisingly, in females, there were no significant differences in DG activity levels relative to controls at any stage of CFM processing (**Fig. 6D**). Importantly, CFM consolidation is highly dependent on post-CFC sleep ^15,19,25^ and DG “engram neuron” reactivation in male mice is sleep-dependent ^15^. However, no significant differences in sleep behavior were observed between male and female experimental consolidation mice (**Fig. 1C**). Together, these data suggest that DG activation patterns during CFM consolidation differ between male and female mice, and that these differences are not attributable to differences in post-CFC sleep.

**Figure 6:**
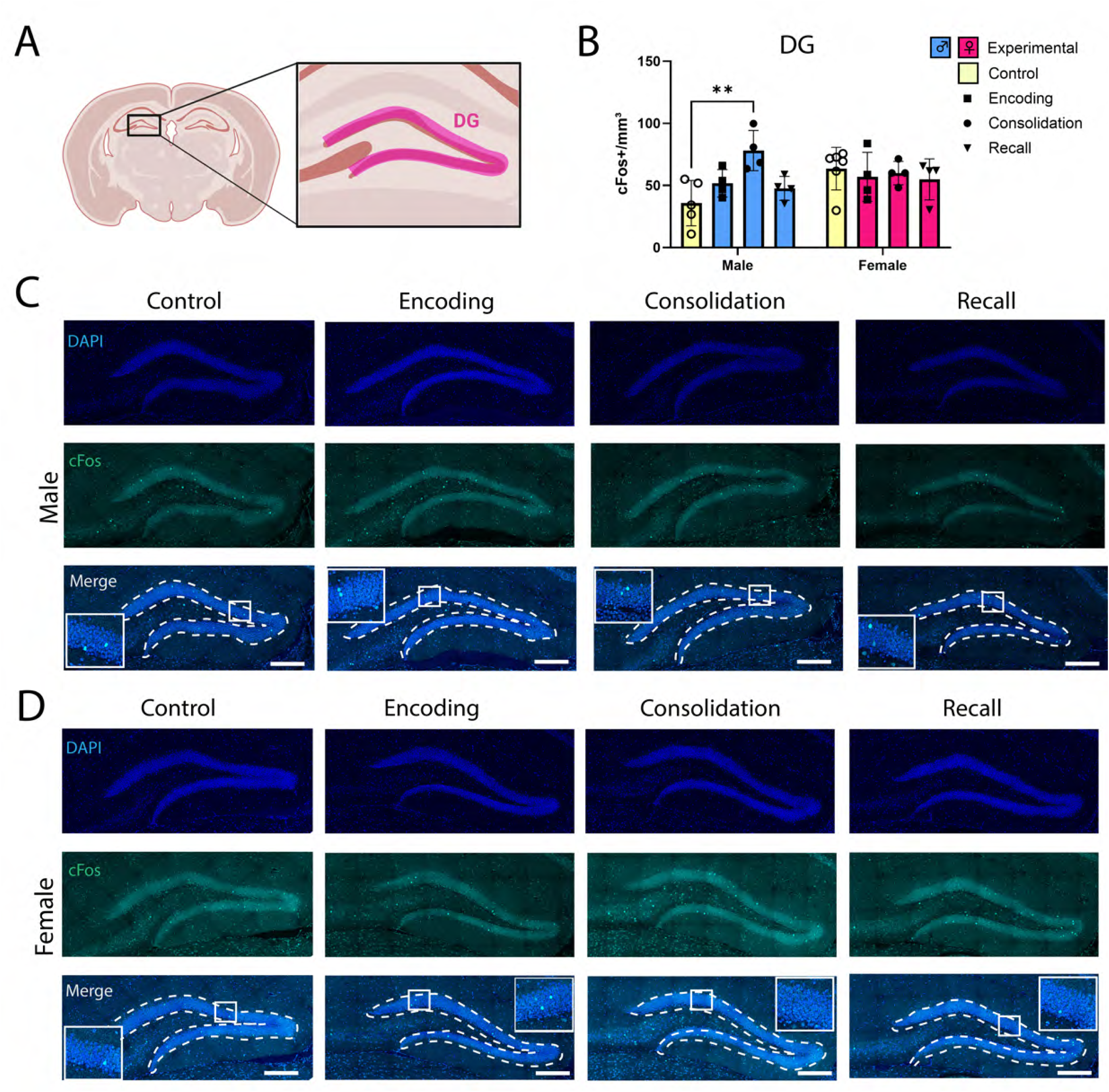
DG granule cell activity increases during CFM consolidation in male mice only. **A)** cFos expression was quantified in the granule cell body layers of dorsal hippocampal DG (inferior and superior blade). **B)** Male mice - but not female mice - showed significantly greater cFos+ neuron density during CFM consolidation compared to controls (two-way ANOVA with Dunnett’s *post hoc* test male consolidation vs. control *p* = 0.0011, all other comparisons *p* > 0.05), with a significant sex × memory stage interaction effect (*n* = 5 male, 6 female control; 4 male, 4 female encoding; 4 male, 4 female consolidation; 4 male, 4 female recall. Main effect of sex: F [1, 27] = 1.08, *p* = 0.31, η^2^ = 0.023; main effect of memory stage: F [3, 27] = 2.77, *p* = 0.061, η^2^ = 0.18; sex × memory stage interaction: F [3, 27] = 3.39, *p* = 0.032; η^2^ = 0.22). **C-D)** Representative DG images from **C)** male and **D)** female mice in each group. Values indicate mean ± SEM. Scale bars = 250 μm.

## Discussion

Our present findings add to a growing body of data suggesting that neural activation patterns differ between male and female mice in some - but not all - brain regions during CFM processing. Regions in which we observed sex differences, and/or trends for sex × memory stage interaction effects, in cFos expression included most major subdivisions of dorsal hippocampus (**Figures 3–6**). However, encoding-associated amygdalar cFos expression was similar between male and female mice, with both increases in cFos during CFM encoding (**Fig. 2B; Supplementary Figure 2**). In all brain regions quantified, we found a significant main effect of memory processing stage (i.e., encoding, consolidation, or recall), but in amygdala there was no significant effect of sex, nor sex × memory stage interaction effects.

There is a growing body of research that suggests there may be sex differences in CFM processing in mice ^29,31^. Recent data suggest biological sex and sex hormones can affect the extent of CFM context generalization and CFM extinction behaviors ^29,54,55^. Moreover, males and females differ with respect to how specific experimental conditions - e.g. single housing or the sex of the experimenter - affect their freezing scores ^41,56^. However, there are also several studies finding no observable sex differences in CFM recall, despite the cellular and molecular mechanisms required for CFM differing between males and females ^37–39^. Similarly, in our present study, no differences in contextual freezing are observed between males and females during either training or testing (**Fig. 1**), yet differences in hippocampal activity patterns during encoding and consolidation are present (**Fig. 3–6**).

One notable difference we observed was the comparatively high activation of both CA1 and CA2 neurons during CFM encoding in females (**Figs. 3 and 5**). While CA1 has been well studied with respect to contextual memory processing ^13,19–21^, much less is known about the contribution of adjacent CA2. While CA2 has synaptic connections with CA1, CA3, DG, and the entorhinal cortex ^57,58^, the region long had a contested role in memory processing due to its unique resistance to synaptic plasticity ^59^. More recently, CA2 has been implicated in social memory processing ^60–62^ and our present findings contribute to a growing body of evidence suggesting it may play a more general role in processing other memory types ^53,63^. We find that CA2, like CA1, shows greater activity during CFM encoding in females. Our data suggest that the extent of activation in these areas during CFC is determined in part by sex.

The most surprising findings from the present study are the apparent differences in dorsal hippocampal activity between males and female mice during CFM consolidation. While there were no apparent sex differences in post-CFC sleep behavior, during consolidation, males showed significantly higher DG cFos expression (**Fig. 6**), and significantly lower CA1 and CA3 cFos expression (**Figs. 3 and 4**), compared to controls – changes not observed in females. This suggests that male mice may downregulate activation of the CA1 and CA3 and upregulate DG activity during the memory consolidation stage. All three regions are implicated in episodic memory formation and recall ^13,15,27,50,64–66^, and recent work from our lab and others suggests that they also play essential roles in CFM consolidation^15,46^. Our prior work - using primarily male mice - has suggested that the success of CFM consolidation is tightly correlated with activation patterns among DG granule cells ^15,46,47^. Both DG granule cell activation and CFM consolidation are strongly affected by post-CFC sleep ^15,26,46,47^. It is unclear whether, and how, activation of DG granule cells leads mechanistically to suppression of CA1 and CA3 pyramidal neuron activity during CFM consolidation in males, although recent data from our lab indicate that patterns of DG interneuron activation can substantially influence activity in both CA structures ^50^. Since CFM consolidation is clearly unimpaired in female mice in our hands, these data suggest that: 1) other circuit-level mechanisms may be involved in the consolidation process in females, and/or 2) differences in encoding between males and females (with encoding evoking stronger activation of CA regions in females) may lead to a stronger memory trace without the need for further hippocampal circuit activation in the hours following CFC. Interestingly, some studies find that males outperform females on context discrimination tasks ^67^. Thus, one possibility is that augmented post-CFC engram neuron reactivation of the DG ^15^, and/or active suppression of pyramidal cells in CA1 and CA3 in males, contributes to this difference by improving pattern separation ^68,69^. Recent data from our lab suggest that this post-CFC reactivation, like CFM consolidation itself ^20,23,25,33^, is sleep-dependent ^15,26,46^. This hypothetical function, linking sex differences in circuit activity to differences in cognition, warrants further investigation.

There are a few important limitations to data interpretation for the present study. First, while some, but not all, studies have identified changes in CFM as a function of the estrus cycle in rodents ^56,70–72^, it is unclear whether, or how, those changes are linked to potential differences in brain activity patterns during CFM processing. While variability of cFos expression measures in female mice does not appear to be different from that of male mice, our present study is not statistically powered to test for differences in cFos expression across the estrus cycle (**Supplementary Table 1**). Thus, future studies will be needed to determine whether cycling hormones affect the activation of different brain regions during CFM processing, and how this might relate to behavioral outcomes. A related limitation is the relatively small sample sizes used in the present studies, which were not powered to detect group differences with small effect sizes. Indeed, for statistically significant group differences reported in these studies, the effect sizes were quite large (i.e., η^2^ > 0.2). For this reason, negative results in these studies should be interpreted with caution. It is also worth noting that while cFos is widely used as an immunohistochemical indicator of prior neuronal activity, it is only one of several such indicators, which can differentially respond to activity within different cell populations ^46,73–75^. Critically, because the expression of other neuronal activity markers may differ between males and females ^76^, we cannot rule out additional sex differences to activity-regulated protein abundance or post-translation modification. In other words, measurement of cFos alone is unlikely to provide a holistic characterization of neuronal or network activity patterns. Lastly, while we did not find significant differences in overall sleep time between male and female mice (**Fig. 1 D-E**), differences in sleep architecture – e.g., differences in the relative proportion of NREM vs. REM sleep – may be present. Future studies using polysomnographic recording may determine whether the observed sex differences in cFos expression during CFM consolidation are driven by more subtle sex differences in post-CFC sleep architecture or brain activity.

It is worth noting parallels between the findings of this study and recent human functional brain imaging studies. In these studies, sex differences have been observed in healthy adults with respect to recall of emotionally valanced episodic and autobiographical memories ^77–79^. Sex differences in memory performance have also been observed in male vs. female patients with mild cognitive impairment or post-traumatic stress disorder ^80,81^. In many of these studies - but not all - differences in performance have been linked to changes in brain activity in the hippocampus, amygdala, and/or neocortical structures during memory encoding and/or recall ^77–79^. As was the case with our present findings, performance on some tasks, such as spatial navigation, is equivalent between men and women, despite apparent sex differences in connectivity between the brain structures they engage ^82^. One important consideration for interpreting our current findings is that men and women have been shown to have different developmental trajectories for wiring connections within the limbic system - both during adolescence ^83^ and in adulthood ^84^. Men and women also differ with respect to age related degeneration and metabolic changes within these structures ^85^. Future studies will be needed to assess how these sex differences over the lifespan affect the dynamic functioning of brain circuitry in the context of healthy cognition. However, data such as our present findings suggest that sex differences in the spatial and temporal circuit dynamics represent different neurobiological processes that achieve the same cognitive outcome.

## Supplementary Materials

**Supplementary Table 1:** Estrus cycle stages of female mice in each experimental group. Estrus cycle stage for 16/18 female mice was assessed visually prior to euthanasia. Stage could not be determined in 2/18 females.

|  | Proestrus (*n* = 1) | Estrus (*n* =7) | Metestrus (*n* =5) | Diestrus (*n* =3) |
| --- | --- | --- | --- | --- |
| Control | 0 | 2 | 2 | 2 |
| Encoding | 0 | 2 | 0 | 1 |
| Consolidation | 1 | 1 | 1 | 0 |
| Recall | 0 | 2 | 2 | 0 |

**Supplementary Figure 1: Locomotor activity patterns during CFC. A)** The average velocity of mice within the testing chamber pre- and post-shock during CFC training was measured in cm/s and compared between males and females. Both male and female mice show significant decreases in velocity following shock administration, consistent with increased post-shock freezing behavior (uncorrected Fisher’s LSD male pre-shock vs. post-shock *p* < 0.0001, female pre-shock vs. post shock *p* < 0.0001). However, the post-shock velocity of female mice was greater than that of male mice (post-shock male vs. female *p* = 0.0096, all other comparisons *p* > 0.05) (*n* = 12 males, 12 females; main effect of sex: F [1, 22] = 4.982, *p* = 0.0361, η^2^ = 0.094; main effect of memory stage: F [1, 22] = 87.83, *p* < 0.0001, η^2^ = 0.38; sex × memory stage interaction: F [1, 22] = 2.58, *p* = 0.12, η^2^ = 0.12). **B)** The average velocity of male and female mice during CFM retrieval was similar (unpaired Student’s t test *p* > 0.05, *n* = 4 males, 4 females, t = 1.31, df = 6, R^2^ = 0.22). **C)** The total distance traveled pre-and post-shock during CFC was measured in cm and compared across sexes. Both male and female mice show significant decreases in distance traveled following shock administration (two-way ANOVA with uncorrected Fisher’s LSD male pre-shock vs. post-shock *p* < 0.0001, female pre-shock vs. post-shock *p* < 0.0001), and there were no significant differences between male and female mice (all other comparisons p > 0.05) (*n* = 12 males, 12 females; main effect of sex: F [1, 22] =2.93, *p* = 0.10, η^2^ = 0.0079; main effect of memory stage: F [1, 22] = 631.6, *p* < 0.0001, η^2^ = 0.90; sex × memory stage interaction: F [1, 22] = 1.04, *p* = 0.32, η^2^ = 0.0015). **C)** The total distance traveled of mice within the testing chamber during CFM retrieval similar between male and female mice (unpaired Student’s t test *p* > 0.05, *n* = 4 males, 4 females, t = 1.31, df = 6, R^2^ = 0.22).

**Supplementary Figure 2: Comparisons of BLA and LA activity patterns during CFM processing. A)** In both males and females, the proportion of cFos+ BLA neurons was significantly increased during CFM encoding (two-way ANOVA with Dunnett’s post hoc test males encoding vs. control *p* < 0.0001, female encoding vs. control *p* = 0.0007). Additionally, males – but not females - showed significantly greater BLA activity during recall compared to controls (male recall vs control *p* = 0.018; all other comparisons *p* > 0.05; main effect of sex F [1, 27] = 0.55, *p* = 0.46, η^2^ = 0.00043; main effect of memory stage F [3, 27] = 25.46, *p* < 0.0001, η^2^ = 0.025; sex × memory stage interaction effect F [3, 27] = 0.032, *p* = 0.99, η^2^ = 0.00044). **B)** Female mice – but not males - had significantly greater LA cFos+ neuron density during encoding compared to controls (two-way ANOVA with Dunnett’s *post hoc* test, female encoding vs. control *p* = 0.013, all other female comparisons *p* > 0.05), while males – but not females – had significantly lower LA cFos+ neuron density during consolidation compared to controls (Dunnett’s *post hoc* test, male consolidation vs. control *p* = 0.0043, all other male comparisons *p* > 0.05). There was a significant main effect of memory stage (main effect of sex F [1, 27] = 0.66, *p* = 0.42, η^2^ = 0.0020; main effect of memory stage F [3, 27] = 12.90, *p* < 0.0001, η^2^ = 0.57; sex × memory stage interaction effect F [3, 27] = 0.44, *p* = 0.72, η^2^ = 0.038).

## Author Contributions

KOM, TY, and AP carried out experiments, collected and analyzed data, KOM and SJA designed experiments and wrote manuscript.

## Animal Use Approval

All mouse procedures were approved by the University of Michigan Institutional Animal Care and Use Committee, under protocol PRO00011982.

## Data Availability Statement

Data used in these studies will be made available upon reasonable request.

## Acknowledgments

The authors are grateful to the members of the Aton lab for helpful commentary on this manuscript, and to Gregg Sobocinski of the MCDB Imaging Core for expert training and assistance with microscopy.

## References cited

1. Beery, A., and Zucker, I. (2011). Sex bias in neuroscience and biomedical research Neurosci Biobehav Rev 35, 565–572.

2. Tronson, N. (2018). Focus on females: A less biased approach for studying strategies and mechanisms of memory. Curr Opin Behav Sci 23, 92–97.

3. Aggarwal, N., and Mielke, M. (2023). Sex Differences in Alzheimer’s Disease. Neurol Clin 41, 343–358.

4. Beam, C., Kaneshiro, C., Jang, J., Reynolds, C., Pedersen, N.L., and Gatz, M. (2018). Differences Between Women and Men in Incidence Rates of Dementia and Alzheimer’s Disease. J Alzheimers Dis 64, 1077–1083.

5. Bourquard, T., Lee, K.H., Al-Ramahi, I., Pham, M., Shapiro, D., Lagisetty, Y., Soleimani, S., Mota, S., Wilhelm, K., Samienasab, M., et al. (2023). Functional variants identify sex-specific genes and pathways in Alzheimer’s Disease. Nat Commun 14, 2765.

6. Eissman, J., Dumitrescu, L., Mahoney, E., Smith, A., Mukherjee, S., Lee, M., Scollard, P., Choi, S., Bush, W.S., Engelman, C.D., et al. (2022). Sex differences in the genetic architecture of cognitive resilience to Alzheimer’s disease. Brain 145, 2541–2554.

7. Mishkin, M., Suzuki, W., Gadian, D., and Vargha-Khadem, F. (1997). Hierarchical organization of cognitive memory. Philos Trans R Soc London B Biol Sci 352, 1461–1467.

8. Fortin, N., Agster, K., and Eichenbaum, H. (2002). Critical role of the hippocampus in memory for sequences of events. Nat Neurosci 5, 458–462.

9. Eichenbaum, H. (1999). The hippocampus and mechanisms of declarative memory Behav Brain Res 103, 123–133.

10. Moscovitch, M., Rosenbaum, R., Gilboa, A., Addis, D., Westmacott, R., Grady, C., McAndrews, M., Levine, B., Black, S., Winocur, G., and Nadel, L. (2005). Functional neuroanatomy of remote episodic, semantic and spatial memory: a unified account based on multiple trace theory. J Anat 207, 35–66.

11. Tovote, P., Fadok, J., and Luthi, A. (2015). Neuronal circuits for fear and anxiety Nat Rev Neurosci 16, 317–331.

12. Kim, W., and Cho, J.-H. (2020). Encoding of contextual fear memory in hippocampal-amygdala circuit. Nat Commun 11, 1382.

13. Daumas, S., Halley, H., Frances, B., and Lassalle, J.M. (2005). Encoding, consolidation, and retrieval of contextual memory: differential involvement of dorsal CA3 and CA1 hippocampal subregions. Learn Mem 12, 375–382.

14. Pereira, L.M., de Castro, C.M., Guerra, L.T.L., Queiroz, T.M., Marques, J.T., and Pereira, G.S. (2019). Hippocampus and Prefrontal Cortex Modulation of Contextual Fear Memory Is Dissociated by Inhibiting De Novo Transcription During Late Consolidation Mol Neurobiol 56, 5507–5519.

15. Wang, L., Park, L., Wu, W., King, D., Vega-Medina, A., Raven, F., Martinez, J., Enseng, A., McDonald, K., Yang, Z., et al. (2024). Sleep-dependent engram reactivation during hippocampal memory consolidation associated with subregion-specific biosynthetic changes. iScience 27, 109408. 10.1016/j.isci.2024.109408

16. Tudor, J.C., Davis, E.J., Peixoto, L., Wimmer, M.E., van Tilborg, E., Park, A.J., Poplawski, S.G., Chung, C.W., Havekes, R., Huang, J., et al. (2016). Sleep deprivation impairs memory by attenuating mTORC1- dependent protein synthesis. Sci Signal 9, ra41.

17. Igaz, L.M., Vianna, M.R., Medina, J.H., and Izquierdo, I. (2002). Two time periods of hippocampal mRNA synthesis are required for memory consolidation of fear-motivated learning. J Neurosci 22, 6781–6789.

18. Bourtchouladze, R., Abel, T., Berman, N., Gordon, R., Lapidus, K., and Kandel, E.R. (1998). Different training procedures recruit either one or two critical periods for contextual memory consolidation, each of which requires protein synthesis and PKA. Learn Mem 5, 365–374.

19. Ognjanovski, N., Broussard, C., Zochowski, M., and Aton, S.J. (2018). Hippocampal Network Oscillations Rescue Memory Consolidation Deficits Caused by Sleep Loss. Cereb. Cortex 28, 3711–3723. 10.1093/cercor/bhy174.

20. Ognjanovski, N., Schaeffer, S., Mofakham, S., Wu, J., Maruyama, D., Zochowski, M., and Aton, S.J. (2017). Parvalbumin-expressing interneurons coordinate hippocampal network dynamics required for memory consolidation. Nature Communications 8, 15039.

21. Ognjanovski, N., Maruyama, D., Lashner, N., Zochowski, M., and Aton, S.J. (2014). CA1 hippocampal network activity changes during sleep-dependent memory consolidation. Front Syst Neurosci 8, 61.

22. Xia, F., Richards, B.A., Tran, M.M., Josselyn, S.A., Takehara-Nishiuchi, K., and Frankland, P.W. (2017). Parvalbumin-positive interneurons mediate neocortical-hippocampal interactions that are necessary for memory consolidation. eLife 6, e27868.

23. Vecsey, C.G., Baillie, G.S., Jaganath, D., Havekes, R., Daniels, A., Wimmer, M., Huang, T., Brown, K.M., Li, X.Y., Descalzi, G., et al. (2009). Sleep deprivation impairs cAMP signalling in the hippocampus. Nature 461, 1122–1125.

24. Havekes, R., Park, A.J., Tudor, J.C., Luczak, V.G., Hansen, R.T., Ferri, S.L., Bruinenberg, V.M., Poplawski, S.G., Day, J.P., Aton, S.J., et al. (2016). Sleep deprivation causes memory deficits by negatively impacting neuronal connectivity in hippocampal area CA1. eLife 5, pii: e13424.

25. Delorme, J., Wang, L., Kodoth, V., Wang, Y., Ma, J., Jiang, S., and Aton, S.J. (2021). Hippocampal neurons’ cytosolic and membrane-bound ribosomal transcript profiles are differentially regulated by learning and subsequent sleep. Proc Natl Acad Sci USA 118.

26. Delorme, J.E., Kodoth, V., and Aton, S.J. (2019). Sleep loss disrupts Arc expression in dentate gyrus neurons. Neurobiol Learn Mem 160, 73–82.

27. Roy, D.S., Park, Y.-G., Kim, M., Zhang, Y., Ogawa, S., DiNapoli, N., Gu, X., Cho, J., Choi, H., Kamentsky, L., et al. (2022). Brain-wide mapping reveals that engrams for a single memory are distributed across multiple brain regions. Nat Commun 13, 1799.

28. Prendergast, B., Onishi, K., and Zucker, I. (2014). Female mice liberated for inclusion in neuroscience and biomedical research. Neurosci Biobehav Rev 40, 1–5.

29. Keiser, A., Turnbull, L., Darian, M., Feldman, D., Song, I., and Tronson, N. (2016). Sex Differences in Context Fear Generalization and Recruitment of Hippocampus and Amygdala during Retrieval. Neuropsychopharmacology 42, 397–407.

30. Nogueira Lotz, F., Knak Guerra, K., Paula Crestani, A., and Alberto Quillfeldt, J. (2025). Multiple context discrimination in adult rats: sex variability and dynamics of time-dependent generalization of an aversive memory Learn Mem *32*, a054081.

31. Gresack, J., Schafe, G., Orr, P., and Frick, K. (2009). Sex differences in contextual fear conditioning are associated with differential ventral hippocampal extracellular signal-regulated kinase activation. Neuroscience 159, 451–467.

32. Samifanni, R., Zhao, M., Cruz-Sanchez, A., Satheesh, A., Mamtaz, U., and Arruda-Carvalho, M. (2021). Developmental emergence of persistent memory for contextual and auditory fear in mice Learn Mem 28, 414–421.

33. Graves, L.A., Heller, E.A., Pack, A.I., and Abel, T. (2003). Sleep deprivation selectively impairs memory consolidation for contextual fear conditioning. Learn. Mem. 10, 168–176.

34. Swift, K.M., Keus, K., Echeverria, C., Cabrera, Y., Jimenez, J., Holloway, J., Clawson, B.C., and Poe, G. (2020). Sex differences within sleep in gonadally intact rats Sleep *43*, zsz289.

35. Mannino, G., Green, T., Murphy, S., Donohue, K., Opp, M., and Rowe, R. (2024). The importance of including both sexes in preclinical sleep studies and analyses. Sci Rep 14, 23622.

36. Choi, J., Kim, S., Fujiyama, T., Miyoshi, C., Park, M., Suzuki-Abe, H., Yanagisawa, M., and Funato, H. (2021). The Role of Reproductive Hormones in Sex Differences in Sleep Homeostasis and Arousal Response in Mice. Front Neurosci 15. 10.3389/fnins.2021.739236

37. Mizuno, I., Matsuda, S., Tohyama, S., and Mizutani, A. (2022). The role of the cannabinoid system in fear memory and extinction in male and female mice Psychoneuroendocrinology 138, 105688.

38. Turner, M., Ball, O., Ray, W., Helm, R., and Jarome, T.J. (2026). Reductions in protein degradation in the retrosplenial cortex regulate contextual fear memory formation in a sex-independent manner Neurobiol Learn Mem 223, 108127.

39. Martin, K.C., Musaus, M., Navabpour, S., Gustin, A., Ray, W., Helm, R., and Jarome, T.J. (2021). Females, but not males, require protein degradation in the hippocampus for contextual fear memory formation Learn Mem 28, 248–253.

40. Tronson, N., and Keiser, A. (2019). A Dynamic Memory Systems Framework for Sex Differences in Fear Memory. Trends Neurosci 42, 680–692.

41. Sakai, M., Yu, Z.-B., Picotin, R., Kasahara, T., Kikuchi, Y., Ono, C., Hino, M., Kunii, Y., Maejima, Y., Shimomura, K., et al. (2025). Experimenters’ sex modulates anxiety-like behavior, contextual fear, and microglial oxytocin transcription in mice. Behav Brain Res 483, 115480.

42. Meenakshi, P., Kumar, S., and Balaji, J. (2021). In vivo imaging of immediate early gene expression dynamics segregates neuronal ensemble of memories of dual events. Mol Brain 14, 102.

43. Barros, V., Mundim, M., Testa Galindo, L., Bittencourt, S., Porcionatto, M., and Mello, L. (2015). The pattern of c-Fos expression and its refractory period in the brain of rats and monkeys. Front Cell Neurosci 9, 72.

44. Onodera, H., Kogure, K., Ono, Y., Igarashi, K., Kiyota, Y., and Nagaoka, A. (1989). Proto-oncogene c-fos is transiently induced in the rat cerebral cortex after forebrain ischemia Neurosci Lett 98, 101–104.

45. Schoenenberger, P., Gerosa, D., and Oertner, T. (2009). Temporal Control of Immediate Early Gene Induction by Light. PLoS One 4, e8185.

46. Delorme, J., Wang, L., Roig Kuhn, F., Kodoth, V., Ma, J., Martinez, J.D., Raven, F., Toth, B.A., Balendran, V., Vega Medina, A., et al. (2021). Sleep loss drives acetylcholine- and somatostatin interneuron-mediated gating of hippocampal activity, to inhibit memory consolidation. Proc Natl Acad Sci USA 118.

47. Martinez, J.D., Brancaleone, W.P., Peterson, K.G., Wilson, L.G., and Aton, S.J. (2023). Atypical hypnotic compound ML297 restores sleep architecture immediately following emotionally valenced learning, to promote memory consolidation and hippocampal network activation during recall Sleep *46*, zsac301.

48. Varol, A., Esen, E., and Kocak, E. (2022). Repeated Collection of Vaginal Smear Causes Stress in Mice. Noro Psikiyatr Ars 59, 325–329.

49. Byers, S., Wiles, M., Dunn, S., and Taft, R. (2012). Mouse Estrous Cycle Identification Tool and Images. PLoS One 7, e35538.

50. Raven, F., Vankampen, A., He, A., and Aton, S. (2026). Dentate gyrus network regulation by somatostatin- and parvalbumin-expressing interneurons differentially impacts hippocampal spatial memory processing. iScience 29, 115067.

51. Prince, T.M., Wimmer, M., Choi, J., Havekes, R., Aton, S., and Abel, T. (2014). Sleep deprivation during a specific 3-hour time window post-training impairs hippocampal synaptic plasticity and memory. Neurobiol Learn Mem 109, 122–130.

52. Yiu, A.P., Mercaldo, V., Yan, C., Richards, B., Rashid, A.J., Hsiang, H.-L., Pressey, J., Mahedevan, V., Tran, M.M., Kushner, S.A., et al. (2014). Neurons Are Recruited to a Memory Trace Based on Relative Neuronal Excitability Immediately before Training. Neuron 83, 722–735.

53. Alexander, G., Riddick, N., McCann, K., Lustberg, D., Moy, S., and Dudek, S.M. (2019). Modulation of CA2 neuronal activity increases behavioral responses to fear conditioning in female mice. Neurobiol Learn Mem 163, 107044.

54. McDermott, C., Liu, D., and Schrader, L. (2012). Role of gonadal hormones in anxiety and fear memory formation and inhibition in male mice. Physiol Behav 105, 1168–1174.

55. McDermott, C., Liu, D., Ade, C., and Schrader, L. (2015). Estradiol replacement enhances fear memory formation, impairs extinction and reduces COMT expression levels in the hippocampus of ovariectomized female mice Neurobiol Learn Mem 118, 167–177.

56. Matsuda, S., Tohyama, S., and Mizutani, A. (2018). Sex differences in the effects of adult short-term isolation rearing on contextual fear memory and extinction. Neurosci Lett 687, 119–123.

57. Kohara, K., Pignatelli, M., Rivest, A., Jung, H.-Y., Kitamura, T., Suh, J., Frank, D., Kajikawa, K., Mise, N., Obata, Y., et al. (2014). Cell type-specific genetic and optogenetic tools reveal hippocampal CA2 circuits Nat Neurosci 17, 269–279.

58. Cui, Z., Gerfen, C., and Young, W.r. (2013). Hypothalamic and other connections with dorsal CA2 area of the mouse hippocampus J Comp Neurol 521, 1844–1866.

59. Zhao, M., Choi, Y.-S., Obrietan, K., and Dudek, S.M. (2007). Synaptic plasticity (and the lack thereof) in hippocampal CA2 neurons J Neurosci 27, 12025–12032.

60. Kassraian, P., Bigler, S., Gilly Suarez, D., Shrotri, N., Barnett, A., Lee, H.-J., Young, W., and Siegelbaum, S. (2024). The hippocampal CA2 region discriminates social threat from social safety Nat Neurosci 27, 2193–2206.

61. Hitti, F., and Siegelbaum, S. (2014). The hippocampal CA2 region is essential for social memory Nature 508, 88–92.

62. Tzakis, N., and Holahan, M. (2019). Social Memory and the Role of the Hippocampal CA2 Region. Front Behav Neurosci 13, 233.

63. Lehr, A., Kumar, A., Tetzlaff, C., Hafting, T., Fyhn, M., and Stober, T. (2021). CA2 beyond social memory: Evidence for a fundamental role in hippocampal information processing. Neurosci Biobehav Rev 126, 398–412.

64. Ramirez, S., Liu, X., Lin, P.A., Suh, J., Pignatelli, M., Redondo, R.L., Ryan, T.J., and Tonegawa, S. (2013). Creating a false memory in the hippocampus. Science 341, 387–391.

65. Liu, X., Ramirez, S., Pang, P.T., Puryear, C.B., Govindarajan, A., Diesseroth, K., and Tonegawa, S. (2012). Optogenetic stimulation of a hippocampal engram activates fear memory recall. Nature 484, 381–385.

66. Stefanelli, T., Bertollini, C., Luscher, C., Muller, D., and Mendez, P. (2016). Hippocampal Somatostatin Interneurons Control the Size of Neuronal Memory Ensembles. Neuron 89, 1074–1085.

67. Keiser, A.A., Turnbull, L.M., Darian, M.A., Feldman, D.E., Song, I., and Tonson, N.C. (2017). Sex Differences in Context Fear Generalization and Recruitment of Hippocampus and Amygdala during Retrieval. Neuropsychopharmacology 42, 397–407.

68. Lee, H., GoodSmith, D., and Knierim, J.J. (2020). Parallel processing streams in the hippocampus. Curr Opin Neurobiol 64, 127–134.

69. Satchell, M., Butel-Fry, E., Zoureddine, Z., Simmons, A., Ognjanovski, N., Aton, S., and Zochowski, M. (2025). Cholinergic modulation of neural networks supports sequential and complementary roles for NREM and REM states in memory consolidation. PLoS Comput Biol 21, e1013097.

70. Rashid, H., and Ahmed, T. (2018). Gender dependent contribution of muscarinic receptors in memory retrieval under sub-chronic stress. Neurosci Lett 681, 6–11.

71. Blair, R., Acca, G., Tsao, B., Stevens, N., Maren, S., and Nagaya, N. (2022). Estrous cycle contributes to state-dependent contextual fear in female rats. Psychoneuroendocrinology 141, 105776.

72. Cossio, R., Carreira, M., Vasquez, C., and Britton, G. (2016). Sex differences and estrous cycle effects on foreground contextual fear conditioning. Physiol Behav 163, 305–311.

73. Gao, Y.-J., and Ji, R.-R. (2009). c-Fos and pERK, which is a better marker for neuronal activation and central sensitization after noxious stimulation and tissue injury? Open Pain J 2, 11–17.

74. Bulthuis, N., Quintana, L., Stackmann, M., and Denny, C. (2025). Immediate-early genes Arc and c-Fos show divergent brain-wide expression following contextual fear conditioning. Communications Biology 8, 1425.

75. Sun, X., Bernstein, M.J., Meng, M., Rao, M., Rao, S., Sorensen, A.T., Yao, L., Zhang, X., Anikeeva, P.O., and Lin, Y. (2020). Functionally Distinct Neuronal Ensembles within the Memory Engram Cell 181, 410–423.

76. Brokling, J., Brunne, B., and Rune, G. (2022). Sex-dependent responsiveness of hippocampal neurons to sex neurosteroids: A role of Arc/Arg3.1 J Neuroendocrinol *34*, e13090.

77. Coenjaerts, M., Trimborn, I., Androvic, B., Stoffel-Wagner, B., Cahill, L., Philipsen, A., Hurlemann, R., and Scheele, D. (2022). Exogenous estradiol and oxytocin modulate sex differences in hippocampal reactivity during the encoding of episodic memories Neuroimage 264, 119689.

78. Young, K., Bellgowan, P., Bodurka, J., and Drevets, W. (2013). Functional neuroimaging of sex differences in autobiographical memory recall Hum Brain Mapp 34, 3320–3332.

79. Mackiewicz, K., Sarinopoulos, I., Cleven, K., and Nitschke, J. (2006). The effect of anticipation and the specificity of sex differences for amygdala and hippocampus function in emotional memory Proc Natl Acad Sci U S A 103, 14200–14205.

80. Sundermann, E., Biegon, A., Rubin, L., Lipton, R., Mowrey, W., Landau, S., Maki, P., and Alzheimer’s Disease Neuroimaging, I. (2016). Better verbal memory in women than men in MCI despite similar levels of hippocampal atrophy Neurology 86, 1368–1376.

81. Shvil, E., Sullivan, G., Schafer, S., Markowitz, J., Campeas, M., Wager, T., Milad, M., and Neria, Y. (2014). Sex differences in extinction recall in posttraumatic stress disorder: a pilot fMRI study Neurobiol Learn Mem 113, 101–108.

82. Sneider, J., Sava, S., Rogowska, J., and Yurtgelun-Todd, D. (2011). A preliminary study of sex differences in brain activation during a spatial navigation task in healthy adults Percept Mot Skills 113, 461–480.

83. Alarcon, G., Cservenka, A., Rudolph, M., Fair, D., and Nagel, B. (2015). Developmental sex differences in resting state functional connectivity of amygdala sub-regions Neuroimage 115, 235–244.

84. Ficek-Tani, B., Horien, C., Ju, S., Xu, W., Li, N., Lacadie, C., Shen, X., Scheinost, D., Constable, T., and Fredericks, C. (2023). Sex differences in default mode network connectivity in healthy aging adults Cereb Cortex 33, 6139–6151.

85. Murphy, D., DeCarli, C., McIntosh, A., Daly, E., Mentis, M., Pietrini, P., Szczepanik, J., Shapiro, M., Grady, C., Horwitz, B., and Rapoport, S. (1996). Sex differences in human brain morphometry and metabolism: an in vivo quantitative magnetic resonance imaging and positron emission tomography study on the effect of aging Arch Gen Psychiatry 53, 585–594.

