## Supplementary figures and images for "Let’s talk about sex: Male and female mice show similar fear memory retention despite hippocampal activity differences during encoding and consolidation"

### Supplementary Figure 1

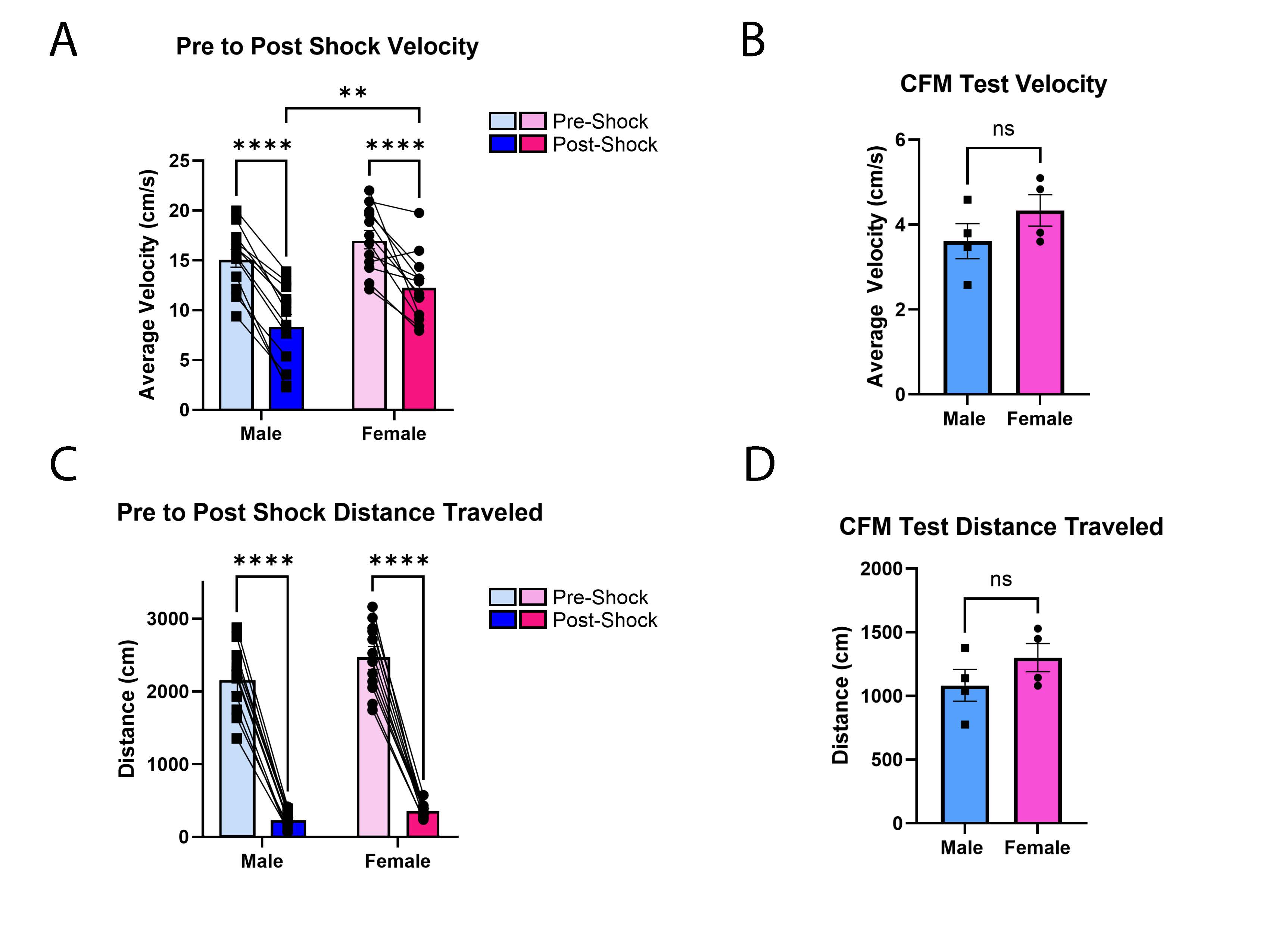

### Supplementary Figure 2

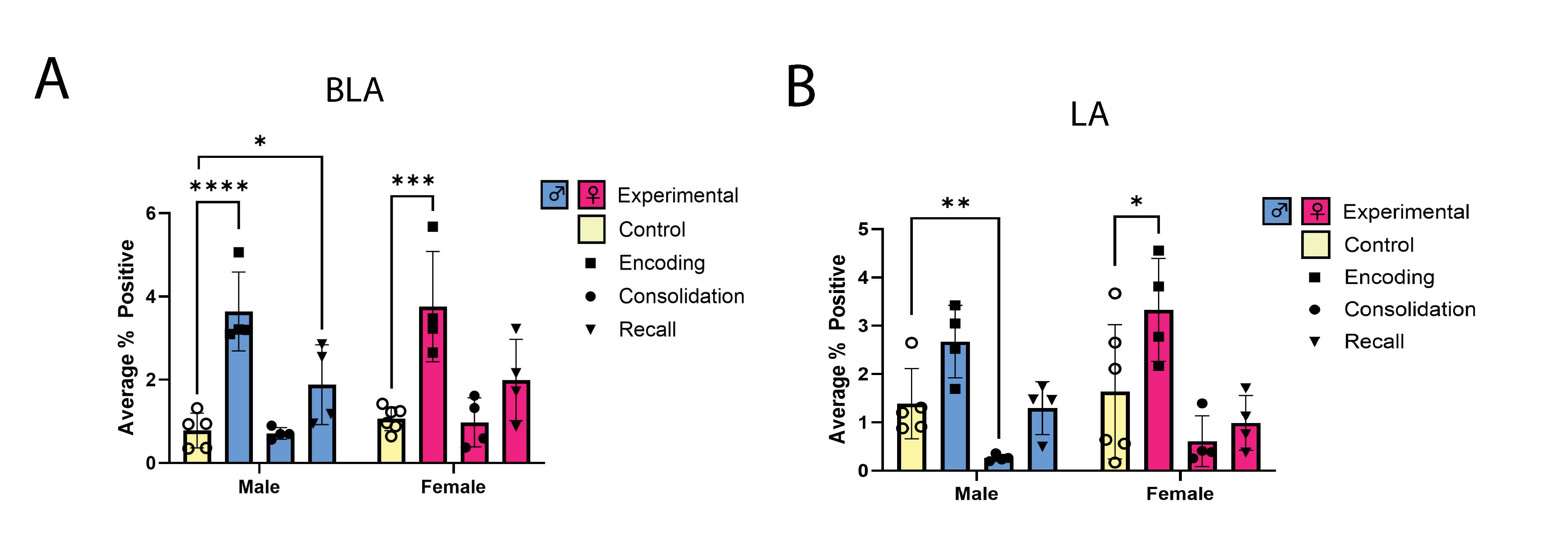
